# Genome-scale comparative analysis for host resistance against sea lice between Atlantic salmon and rainbow trout

**DOI:** 10.1101/624031

**Authors:** Pablo Cáceres, Agustín Barría, Kris A. Christensen, Liane N. Bassini, Katharina Correa, Jean P. Lhorente, José M. Yáñez

**Affiliations:** Facultad de Ciencias Veterinarias y Pecuarias, Universidad de Chile, Av. Santa Rosa 11735, La Pintana, Santiago 8820808, Chile; Fisheries and Oceans Canada 4160 Marine Drive, West Vancouver, BC, Canada; Escuela de Medicina Veterinaria, Facultad de Ciencias de la Vida, Universidad Andres Bello, Santiago, Chile; Benchmark Genetics Chile, Ruta 7 Carretera Austral Km 35, Puerto Montt, Chile; Núcleo Milenio INVASAL, Concepción Chile

**Keywords:** *Caligus rogercresseyi*, *Salmo salar*, *Oncorhynchus mykiss*, GWAS, Parasite Resistance, Comparative Genomics

## Abstract

Sea lice (*Caligus rogercresseyi*) are ectoparasites that cause major production losses in the salmon aquaculture industry worldwide. Atlantic salmon (*Salmo salar*) and rainbow trout (*Oncorhynchus mykiss*) are two of the most susceptible salmonid species to sea lice infestation. The goal of this study was to identify common candidate genes involved in resistance against sea lice. For this, 2,626 Atlantic salmon and 2,643 rainbow trout from breeding populations were challenged with sea lice and genotyped with a 50k and 57k SNP panel. We ran two independent genome-wide association studies for sea lice resistance on each species and identified 7 and 13 windows explaining 3% and 2.7% respectively the genetic variance. Heritabilities were observed with values of 0.19 for salmon and 0.08 for trout. We identified genes associated with immune responses, cytoskeletal factors and cell migration. We found 15 orthogroups which allowed us to identify *dust8* and *dust10* as candidate genes in orthogroup 13. This suggests that similar mechanisms can regulate resistance in different species; however, they most likely do not share the same standing variation within the genomic regions and genes that regulate resistance. Our results provide further knowledge and may help establish better control for sea lice in fish populations.

## Introduction

Sea lice, *Caligus rogercresseyi*, first described in 1997 by Boxshall & Bravo^1^ are currently the most harmful parasite in salmon farming worldwide^2^, and are also the main parasite affecting Chilean salmon production. To date, the economic losses due to this parasite are mainly associated with the reduction of feed conversion and fish growth, indirect mortality, loss of product value and treatment costs. It has been estimated that the global costs for the control of this parasite has reached $436 million (USD) annually^3^. The parasite life cycle is comprised of eight stages of development^4^: two states of nauplii, one copepod state, four chalimus states and the adult state. The stages of nauplii (n1-2) and the stage of copepods (infectious stage) are planktonic stages. The four stages of chalimus (1-4) are sessile stages and the adult is a mobile stage^5^.

It has been observed that *Caligus* primarily affects Atlantic salmon (*Salmo salar* and rainbow trout (*Oncorhynchus mykiss*), while coho salmon (*Oncorhynchus kisutch*) has an innate lower susceptibility to the parasite^6^. The clinical signs attributed to infection by sea lice include skin lesions, osmotic imbalance and greater susceptibility to bacterial and viral infections through the suppression of immune responses by damage to the skin of the host^7^.

Recent studies have estimated significant low to moderate genetic variation for resistance to *C. rogercresseyi*, with heritability values ranging between 0.12 and 0.32 in Atlantic salmon when resistance is defined as the number of parasites fixed to all the fins^5,7,8^. Similarly, when sea lice resistance is defined as the logarithm of the parasite load density, heritability values range between 0.13 and 0.33 in Atlantic salmon^9,10^. Although significant, a heritability value of 0.09 for sea lice resistance have been recently reported in rainbow trout (Bassini et al., submitted)^11^. These results support the feasibility of including resistance to sea lice into breeding programs for both Atlantic salmon and rainbow trout^5,8,12^.

Due to increased research efforts focusing on salmonid genomics, there has been a rapid increase in the discovery of Atlantic salmon and rainbow trout SNP markers and high-density SNP genotyping panels have been developed for both species^13–15^. The availability of a reference genome for rainbow trout^16^ and Atlantic salmon^17^ have allowed genomic regions associated with resistance to diseases to be identified and annotated. For example, it has been possible to identify regions associated with resistance to bacterial diseases such as *Piscirickettsia salmonis*^18^, *Flavobacterium psychrophilum*^19,20^ and also parasitic diseases^7,10^ in both species.

Comparative genomic approaches^21^ allow the identification of genomic similarities between evolutionarily distant species, such as: conserved genes, traces of genome duplication and the comparison of gene function in different species^22^. Traditionally, comparative genomics analyses have focused on orthologous genes (genes related to each other resulting from direct transmission from a common ancestor)^22^. Comparative genomic studies between salmonids have mainly focused on finding evolutionary similarities, such as quantitative trait loci (QTL) related to growth or sexual differentiation^23–25^. To date, there have been no studies comparing genomic regions associated with resistance to *Caligus* in salmonid species.

The objective of the present study was to: i) identify genomic regions associated with resistance to *Caligus rogercresseyi* in Atlantic salmon and rainbow trout through GWAS, and ii) identify functional candidate genes potentially related to trait variation through a comparative genomics approach based on exploring orthologous genes within the associated regions across species.

## Results and Discussion

The comparative genomics analysis used in this study, allowed us to identify groups of ortholog genes and several candidate genes among adjacent SNP that explained more than 1% of the significant genetic variance for resistance to *Caligus rogercresseyi*. This first study which compared resistance in both salmonid species, focused on candidate genes which function either to directly defend against the parasite or that participate in defense mechanisms against this parasite.

There was no difference between the average number of sea lice found on Atlantic salmon or rainbow trout in the experimental challenge (Table 1). An average of 5.9 ± 6.6 and 6.1 ± 4.2 was estimated for Atlantic salmon and rainbow trout, respectively. In terms of the maximum number of parasites, this value varied from 106 parasites in Atlantic salmon to 28 in rainbow trout. For both species, there were animals that did not present with any parasites. The average weight at the end of the experimental challenge was 278.1 ± 90.3 g (ranging from 104 to 569 g) and 173.1 ± 31.4 g (ranging from 86 to 265 g), for Atlantic salmon and rainbow trout, respectively. Although the average number of parasites was not significantly different between the two species, the difference in the average final weight at the end of each challenge could explain the difference in the maximum number of parasites found (~4 times more parasites in the larger Atlantic salmon). The number of parasites counted in each species after the challenge is below the range determined in previous studies. For instance, Ødegård *et al.* (2014)^9^ obtained an average of 20.96 ± 19.68, while Robledo *et al.* (2018)^26^ reported an average of 38 ± 16.

**Table 1.** Summary statistics for body weight (BW) and lice count (LC) in Atlantic salmon and rainbow trout.

| Species | Mean BW<br>(g) | SD<sup>1</sup> BW<br>(g) | Min<br>BW(g) | Max BW<br>(g) | LC mean | LC SD | LC min | LC max |
| --- | --- | --- | --- | --- | --- | --- | --- | --- |
| *S. salar* | 278.1 | 90.3 | 104.0 | 569.0 | 5.9 | 6.61 | 0 | 106 |
| *O. mykiss* | 173.1 | 31.4 | 86.0 | 265 | 6.1 | 4.22 | 0 | 28 |

<sup>1</sup> SD (standard deviation)

To measure resistance to *C. royercresseyi* we used the logarithm of lice density (LogLD), which allows for correction to the number of parasites based on the body weight of each fish^9^. The empirical LogLD distribution for both species is shown in Figure 1. The range of LogLD for Atlantic salmon was greater than for rainbow trout, varying from −4.18 to 1.023 and from −3.69 to 0.03, respectively. The average LogLD distribution in the present study was 2.12 ± 0.89 and 1.64 ± 0.65 for Atlantic salmon and rainbow trout, respectively, which are similar values to those reported in a previous study in a different Atlantic salmon population (between −1.66 ± 0.73 and −2.55 ± 0.58)^9^.

**Figure 1.**
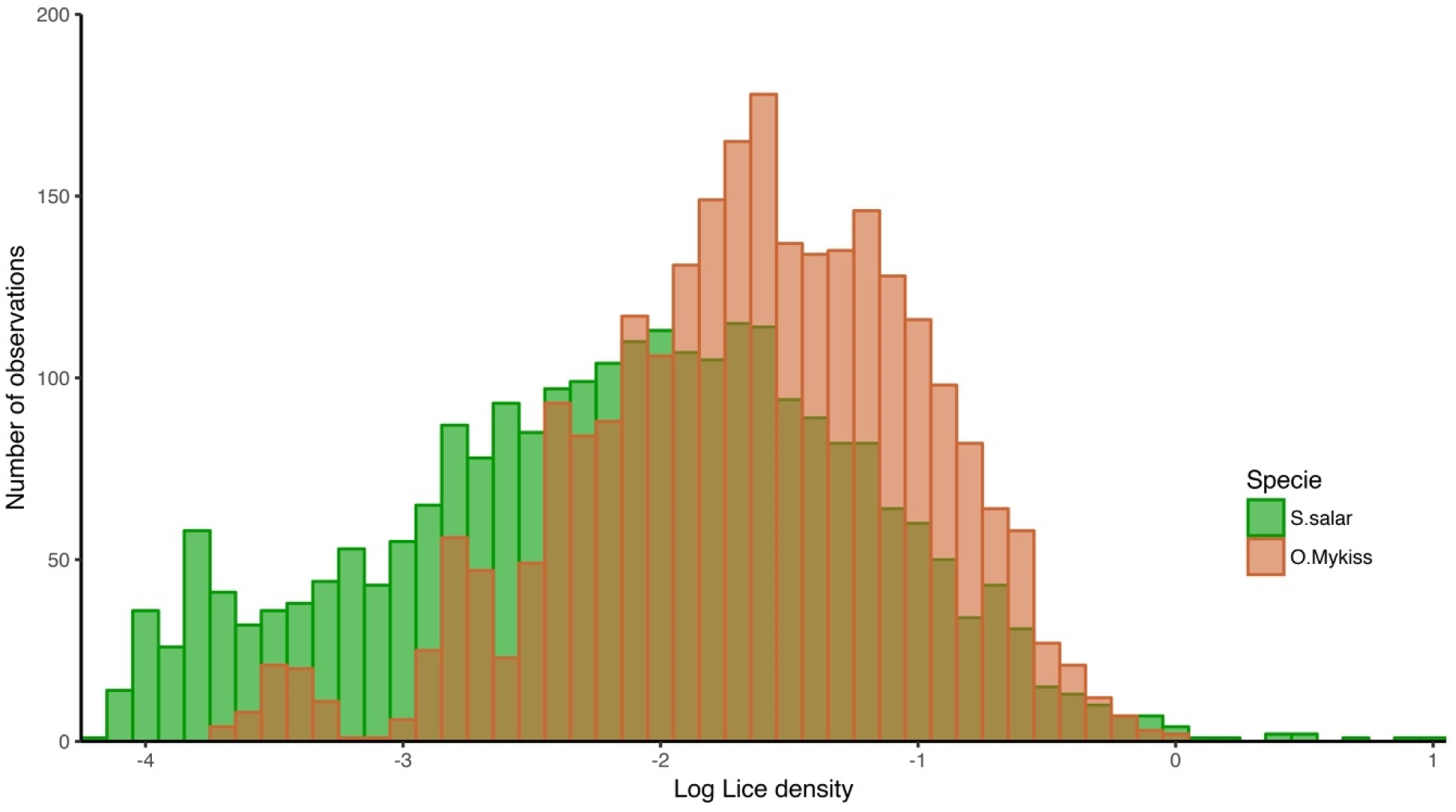
Histogram for log lice density (LogLD) for S. *salar* (green) and *O. mykiss* (orange).

A total of 2,040 (77.6%) Atlantic salmon, and 45,117 (96.7%) SNPs passed genotyping quality control, while in rainbow trout, 2,466 (93.3%) individuals and 27,146 (67.4%) SNP remained. In both species, significant genetic variation for resistance to *C. rogercresseyi* was estimated by using genomic information, with heritability values of 0.19 ± 0.03 and 0.08 ± 0.01 for Atlantic salmon and rainbow trout, respectively (Table 2). Tsai *et al.* (2016)^10^, estimated genomic heritability values for resistance against *L. salmonis* of 0.22 ± 0.08 and 0.33 ± 0.08, while Ødegård *et al.* (2014)^9^, observed heritability values of 0.14 ± 0.03 and 0.13 ± 0.03 in Atlantic salmon. Similarly, Yañez *et al* (2014) and Correa *et al* (2017)^8,12^ estimated values ranging from 0.10 to 0.12 when defining resistance as the total number of parasites found on all fins using genomic and pedigree information, and Lhorente et al (2012)^5^ estimated heritability values of 0.22 ± 0.06 in Atlantic salmon with traits of total count of sessile sea lice per fish and body weight.

**Table 2.** Window position and genetic variance of representative SNP.

| Window Position (bp)<sup>1</sup> | Chr<sup>2</sup> | First and last SNP of window | Var (%)<sup>3</sup> |
| --- | --- | --- | --- |
| **Atlantic Salmon** |  |  |  |
| 111398941-112068765 | 9 | Affx-93376378-Affx-93407247 | 3.00 |
| 46812870-48951308 | 6 | Affx-93402585-Affx-93349216 | 1.97 |
| 38862311-39791664 | 9 | Affx-93301094-Affx-93369286 | 1.76 |
| 21387020-22266613 | 3 | Affx-93351937-Affx-93441285 | 1.48 |
| 13302321-13992142 | 25 | Affx-93449253-Affx-93379811 | 1.33 |
| 53395987-54096750 | 3 | Affx-93397375-Affx-93327854 | 1.08 |
| 47471286-48390124 | 20 | Affx-93269128-Affx-93365186 | 1.05 |
| **Rainbow trout** |  |  |  |
| 55155679-55750071 | 15 | Affx-88957834-Affx-88937607 | 2.77 |
| 14742278- 15862261 | 26 | Affx-88926342-Affx-88913419 | 2.35 |
| 35337013- 36760480 | 28 | Affx-88961757-Affx-88909402 | 1.77 |
| 57059237- 57975684 | 3 | Affx-88911532-Affx-88917594 | 1.73 |
| 14313209- 15091943 | 9 | Affx-88944322-Affx-88927798 | 1.67 |
| 10428810- 11808090 | 2 | Affx-88922063-Affx-88937734 | 1.66 |
| 62615440- 63610148 | 14 | Affx-88942436-Affx-88960365 | 1.43 |
| 27521395-28664751 | 21 | Affx-88932888-Affx-88952962 | 1.31 |
| 43708499-44788897 | 10 | Affx-88912735-Affx-88908041 | 1.29 |
| 26370287-27397531 | 29 | Affx-88925775-Affx-88920967 | 1.25 |
| 50072755-50809742 | 7 | Affx-88956194-Affx-88906442 | 1.17 |
| 15571716-16056405 | 16 | Affx-88940047-Affx-88947010 | 1.07 |
| 7055909-8035171 | 4 | Affx-88935786-Affx-88957720 | 1.03 |

<sup>1</sup>Window position in base pair (BP).
<sup>2</sup>Chromosome (Chr).
<sup>3</sup>Variance (Var)

We found 5 chromosomes harboring 7 loci explaining more than 1% of the genetic variance for sea lice resistance in Atlantic salmon (Figure 2). In general, these regions explained a low percentage of the total genetic variation with a maximum of 3% explained by a single locus. Thus, two SNP windows (a window was defined as 20 contiguous SNPs) in *Ssa3* explained 1% and 1.4% of the genetic variance. In *Ssa6* there was a window that explained up to 1.9% of the genetic variance, while two windows that explained 1.7% and 3% were found in *Ssa9*. In addition, in *Ssa20* and *Ssa25* we found windows that explained 1.05% and 1.33%, respectively. Table 3 shows the variance explained by each window of SNP in both species.

**Figure 2.**
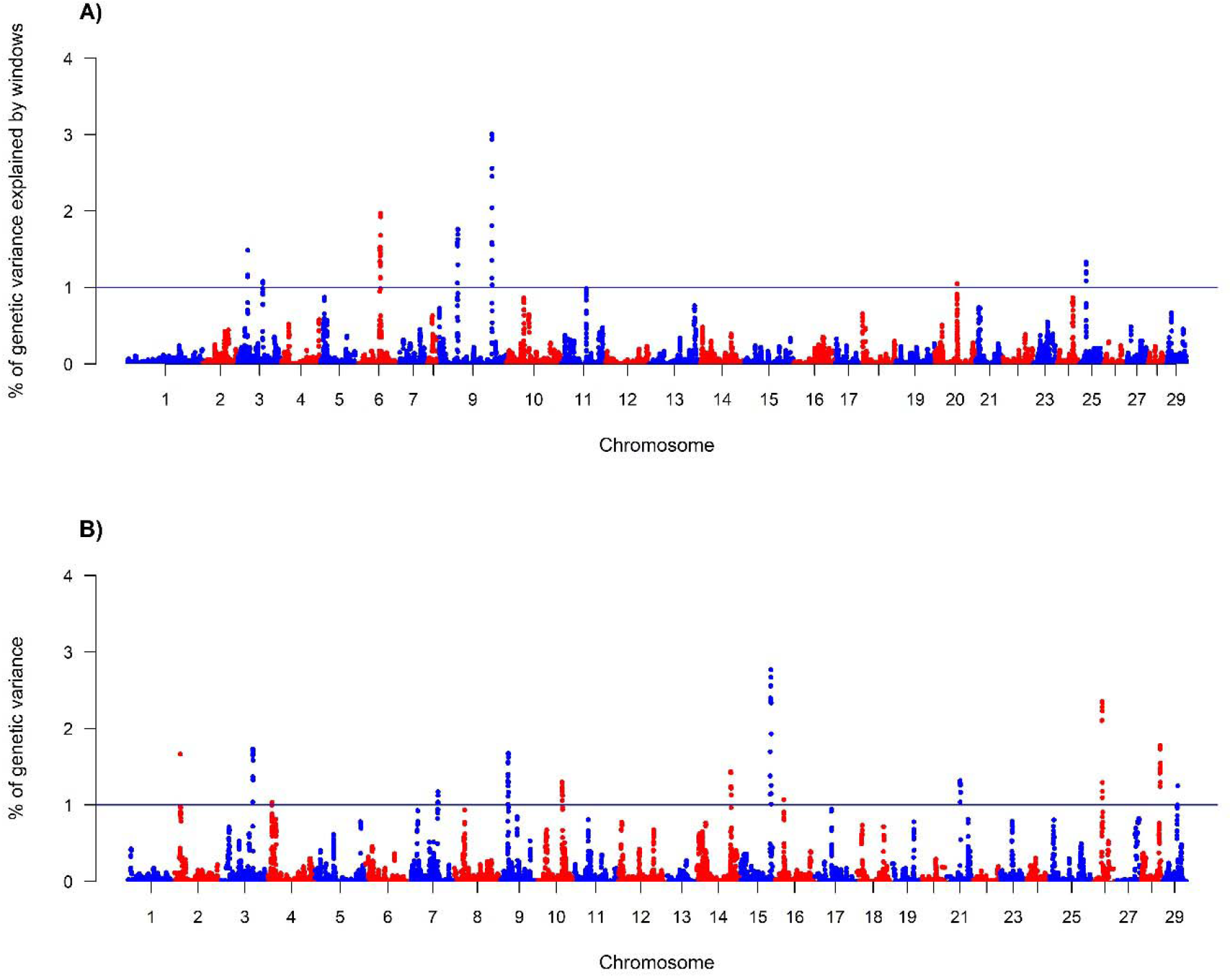
Weighted single-step GBLUP (wssGBLUP) for Log sea lice density (LogLD) in Atlantic salmon (A) and rainbow trout (B). Blue line indicates greater than 1% of the variance explained.

**Table 3.** Genes identified as possible candidates for sea lice resistance in Atlantic salmon.

| Protein name | Gene ID<sup>1</sup> | Location (BP)<sup>2</sup> | Chr<sup>3</sup> | Function |
| --- | --- | --- | --- | --- |
| *Tenascin R* | *tnr* | 22.113,558 - 22.312,716 | 3 | Immune response |
| *T-cell activation Rho GTPase-activating protein* | *tagap* | 47.344,667 - 47.350,357 | 6 | Immune response |
| *Forkhead box protein N1-like isoform X1* | *LOC106612922* | 111.631,268 - 111.668,978 | 9 | Immune response |
| *Immunoglobulin superfamily member 11-like* | *igsf11* | 47.453,435 - 47.570,948 | 20 | Immune response |
| *Tripartite motif-containing 45* | *trim45* | 31.699,662 - 31.705,320 | 25 | Immune response |
| *pre-B-cell leukemia transcription factor 1-like* | *LOC106607421* | 43.977,204 - 44.037,235 | 6 | Immune response |
| *Bromodomain-containing protein 4-like isoform* | *LOC106600922* | 53.482,098 - 53.526,053 | 3 | Immune response |
| *PAPPALYSIN-2* | *pappa2* | 21.972,287 - 22.050,408 | 3 | Metalloprotease |
| *Carboxypeptidase D* | *cdpa* | 111.778,091 - 111.828,613 | 9 | Metalloprotease |
| *GEM-interacting protein-like isoform X3* | *LOC106600913* | 53.672,635 - 53.709,985 | 3 | Metalloprotease |
| *Epidermal growth factor* | *egf* | 55.552,926 - 55.584,698 | 9 | Cytoskeletal |
| *Procollagen galactosyltransferase 1* | *LOC106607427* | 43.820,342 - 43.851,337 | 6 | Cytoskeletal |
| *Heat shock protein HSP 90-beta* | *hs90b* | 48.362,362 - 48.374,359 | 6 | Cytoskeletal |
| *Collagen alpha-1(XXVIII) chain-like* | *col28a1* | 13.872,240 - 13.923,778 | 25 | Cytoskeletal |
| *gap junction alpha-4 protein-like* | *LOC106611294* | 39.373,024 - 39.373,931 | 9 | Cytoskeletal |
| *Pleckstrin homology domain-containing family H member 1-like* | *LOC106611291* | 39.101,628 - 39.165,084 | 9 | Cytoskeletal |
| *serine/threonine-protein kinase OSR1-like* | *LOC106600551* | 44.726,186 - 44.781,350 | 3 | Cytoskeletal |
| *rho-related GTP-binding protein RhoB-like* | *LOC106607565* | 46.401,032 - 46.403,034 | 6 | Cytoskeletal |
| *pleckstrin homology and RhoGEF domain containing G1* | *plekhg1* | 64.024,693 - 64.178,709 |  | Cytoskeletal |

<sup>1</sup>Gene identification (Gene ID) from the NCBI database GenBank assembly accession:
GCA\_000233375.4.
<sup>2</sup>Location in base pair (BP).
<sup>3</sup>Chromosome (Chr).

For rainbow trout the GWAS for LogLD identified 13 regions located in 13 different chromosomes that exceeded 1% of the total genetic variance (Figure 2). Similar to Atlantic salmon, these windows explained a low percentage of the total variance with a maximum of 2.77%; nevertheless, the number of regions surpassing 1% of the genetic variance explained was almost double in rainbow trout compared to Atlantic salmon. The important genomic regions in rainbow trout were located on chromosomes *Omy2, Omy3, Omy4, Omy7, Omy9, Omy10, Omy14, Omy15, Omy16, Omy21, Omy26, Omy28 and Omy29*, and explained 1.6%, 1.7%, 1.03%, 1.17%, 1.6%, 1.2%, 1.4%, 2.77%, 1.07%, 1.3%, 2.3%, 1.7% and 1.2% of the genetic variance for LogLD, respectively (Table 3).

As has been previously reported for other disease resistance traits in aquaculture species^18,20,27–29^, our results suggest that sea lice resistance is mainly of polygenic nature (i.e. many genes with small effect are involved in the trait). These results agree with previous studies on sea lice resistance, where a similar genetic architecture was suggested by Tsai *et al.* (2016), Rochus *et al* (2018) and Correa *et al.* (2017)^7,10,30^ for *Lepeophtheirus salmonis* and *C. royercresseyi* resistance. Recently Robledo et al (2018)^31^ described three QTL in Atlantic salmon related to sea lice resistance, using RNA-seq and WGS data. Since sea lice resistance is polygenic, the genetic improvement of sea lice resistance would most likely be best accomplished by means of genomic selection instead of marker assisted selection. For instance, Correa *et al* (2017) and Tsai *et al* (2016)^10,12^ have shown an increase in the accuracy of estimated breeding values (EBVs) using genomic selection, over the use of pedigree-based models^9^.

The exploration of the genes within the windows that explained over 1% of the genetic variance for LogLD showed a series of possible candidate genes that were classified into three groups: related to the immune response, cytoskeleton or metalloproteases. The genes are listed in Table 3 and Table 4 for Atlantic salmon and rainbow trout, respectively.

**Table 4.**
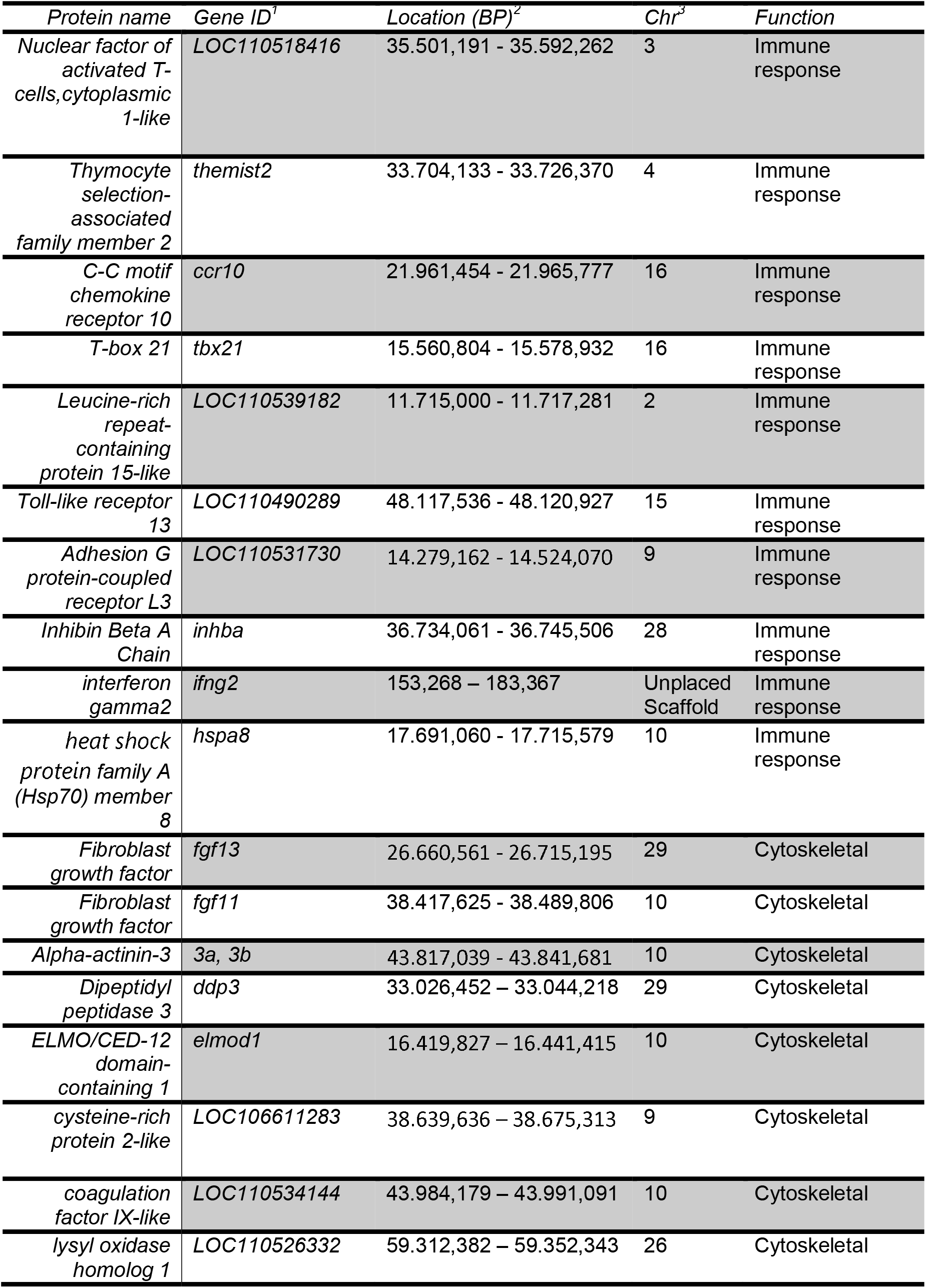
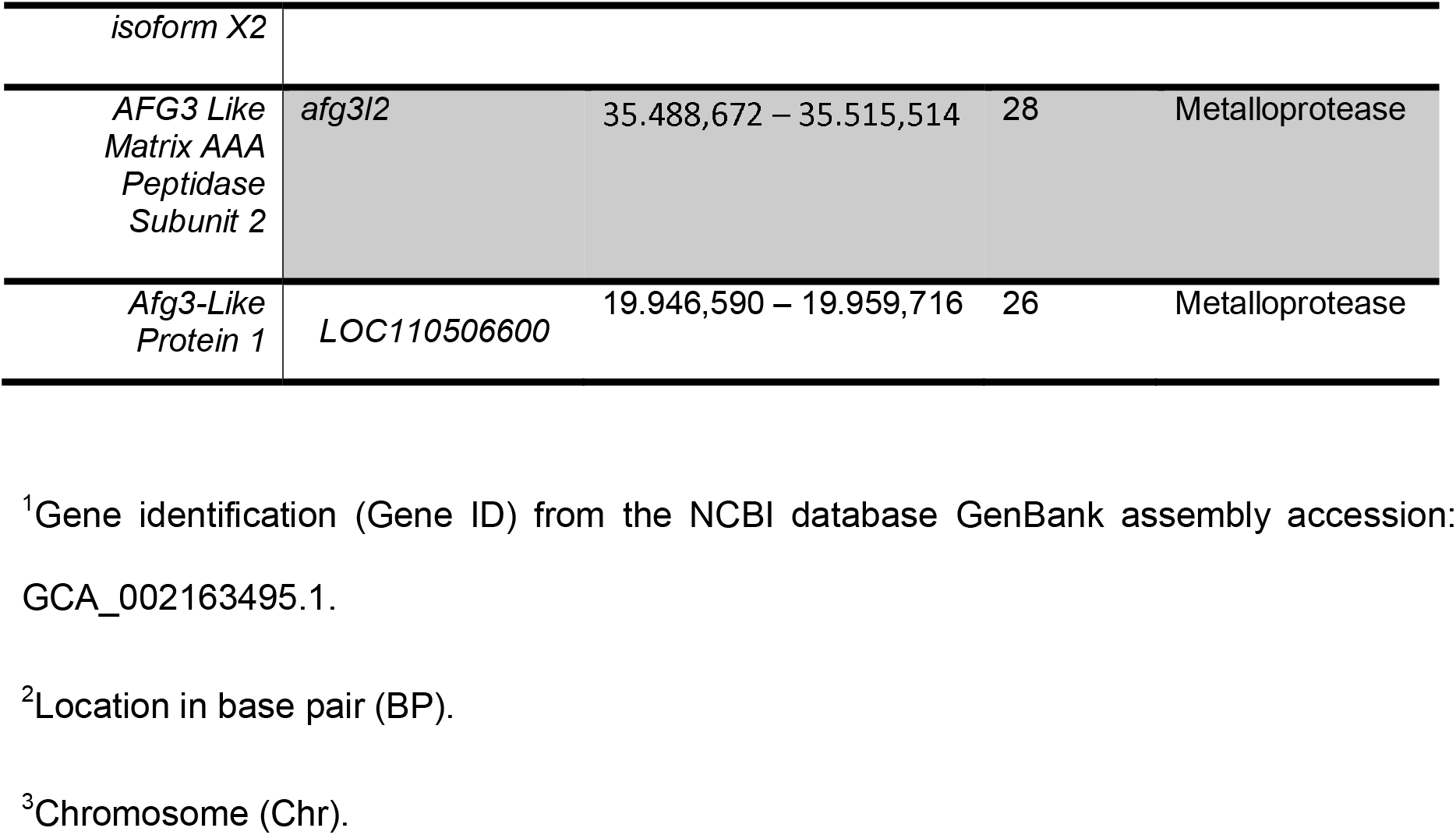
Genes identified as possible candidates for sea lice resistance in Rainbow trout.

In salmonids, the main response of the immune system to parasites is mediated by T-Helper 1 and T-Helper 2^32^ cells. Thus, genes related with immune response, either by promoting leukocyte growth or favoring migration or activation are strong candidate genes. For instance, in Atlantic salmon we found *T-cell activation Rho GTPase-activating protein* (TAGAP) which participates in the activation and recruitment of T cells by cytokines^33^, and *tenascin R* (TNR) which is an extracellular matrix protein, present in bone marrow, thymus, spleen and lymph nodes^34^. The latter has been described as having an adhesin function favoring the mobility of lymphocytes and lymphoblasts^34,35^. In rainbow trout, we found candidate genes with similar functions, such as, T-box 21 (*tbx21*), also known as T-bet (*T-box expressed in T cells*). This gene belongs to the sub Tbr1 family^36^ and generates type 1 immunity and participates in the maturation and migration of T-helper 1 (Th1) cells, which in turn produce interferon-gamma (IFN-γ). Studies have described T-bet expression in NK cells (natural killer), dendritic cells and T CD8+ cells^37,38^. A recent study^26^ on gene expression with *C. rogercresseyi* infestation in susceptible and resistant Atlantic salmon indicated that several components of the immune system (inflammatory response, cytokine production, TNF and NF-kappa B signaling and complement activation) and tissue repair are upregulated during infection.

*Forkhead box protein N1-like* (FOXN1) present on *Ssa9* of Atlantic salmon is part of a family of genes widely studied in humans, which are related to various functions including cell growth, lymph node development and T cell differentiation^39^. In addition, it has been proposed that FOXN1 has a role in the activation of fibroblast growth factor receptors^39^.

Meanwhile in trout on *Omy21, serine/threonine-protein phosphatase 2A 56 kDa* was identified, which is described as having participated in cell growth and signaling^40^. Robledo *et al.* (2018)^26^ recently found that in Atlantic salmon, this protein showed the most significant change in the expression ratio between healthy skin and skin where sea lice were found^26^. In Atlantic salmon, we identified *Tripartite motif-containing protein 45* (TRIM) on *Ssa25* which belongs to a large family of proteins present in diverse organisms that can function as a ligase, and can modify ubiquitins and proteins stimulated by interferon of 15 kDa (ISG15)^41^.

Warm-water fish, such as Zebrafish (*Danio rerio*) and rita catfish (*Rita rita*), lower infiltration of neutrophils, favoring wound closing by means of accelerated growth of the epidermis which can take place in a few hours^42^. In coho salmon (*Oncorhynchus kisutch*) it has been observed that a neutrophil infiltration occurs until the second day after sea lice infestation, together with an inflammatory reaction and hyperplasia in the zone^43^, with posterior leukocyte recruitment and migration.

Several metalloproteases were found in both species, but for the interest of this study, we focused on GEM-interacting protein which interacts with Rab27a or its effector in leucocytes. Rab is a large family of small GTPasas responsible for vesicle cellular transport^44^. Deficiencies of this molecule or the related human protein (?), is correlated with immune deficiencies due to the malfunction of cytotoxic activity of T-lymphocytes, natural killer cells and neutrophils^45^.

Considering the importance of cell growth and movement in response to sea lice infestation, the cytoskeleton may play a considerable role in this response as well. Genes related to the cytoskeleton were identified, such as epidermal growth factor (EGF), found on *Ssa9*. This gene is part of a superfamily of receptors with tyrosine kinase activity that have been described in a variety of organs with growth promoter functions, cellular differentiation^46^ and could participate in tissue repair by promoting cell growth^36^. In rainbow trout, fibroblast growth factors (*fgf11 - Omy10, fgf13 - Omy29*) have similar functions (angiogenesis and pro-inflammatory response), and were identified as important genes involved in sea lice resistance by Skugor *et al.* (2009) and Robledo *et al.* (2018) in Atlantic salmon^26,43^

*ELMO/CED-12 domain-containing prot 1* was identified on *Omy10* of the trout. This protein is characterized mainly from research in the model organism *C. elegans. ELMO/CED-12 domain-containing prot 1* participates in the phagocytosis of apoptotic cells, and in mammals it also has a role in cell migration^47^. Other cytoskeleton related candidate genes include: *Procollagen galactosyltransferase 1* present on *Ssa6, collagen alpha-1 (XXVIII) chain-like* on *Ssa25* and *pleckstrin homology domain-containing family H member 1-like*^48^.

The top ten SNPs that explained the greatest variance are located on *Ssa9* of the Atlantic salmon, in close proximity to the *breast carcinoma-amplified sequence 3 (bcas3)* gene, which in Atlantic salmon codes for a cell migration factor associated with microtubules that favors cellular mobility^49^. Cell migration is generally induced in response to chemotactic signals, which induces changes in the cytoskeleton and extracellular matrix^50^.

We also found in trout, the *tripartite motif-containing protein 16-like* on *Omy15*, which is part of the TRIM superfamily and has functions related to cell differentiation, apoptosis, regulation of transcription and signaling pathways^41^ This gene is similar to *Tripartite motif-containing protein 45* present on *Ssa25*. In this region, we also found a locus that codes for interferon-γ 2 (*ifng2*), which is a cytokine that participates in type 1 immune responses and that favors the presentation of antigens and activation of macrophages^51^. On this same chromosome (*Omy15*), we also identified *putative ferric-chelate reductase 1 (frrs1)*, which has been described as having functions in the fixation of iron in teleosts^52^. Robledo *et al.* (2018)^26^ identified *heme-binding protein* 2 (HEBP2) as a gene involved in Atlantic salmon sea lice resistance, which has an iron-binding function. Different authors^53,54^ have stated that decreasing the availability of iron can be part of a nutritional defense mechanism against sea lice infestation.

The analyses performed show regions of synteny between both species (Figure 3): there are homologous regions that share similarity between the chromosomes across species. However, there was no obvious shared sea lice resistance associations between Atlantic salmon and rainbow trout (Figure 3). It is possible that similar mechanisms regulate resistance between the two species, but the examined populations did not share the same standing variation of the genes regulating resistance. For example, *Ssa03* (Atlantic salmon) shares homology with *Omy28* (rainbow trout) and *Ssa25* with *Omy03*. When performing the search for genes by chromosome (see Table 4), it was not possible to identify genes that were shared in the indicated regions that were related to the trait studied.

**Figure 3.**
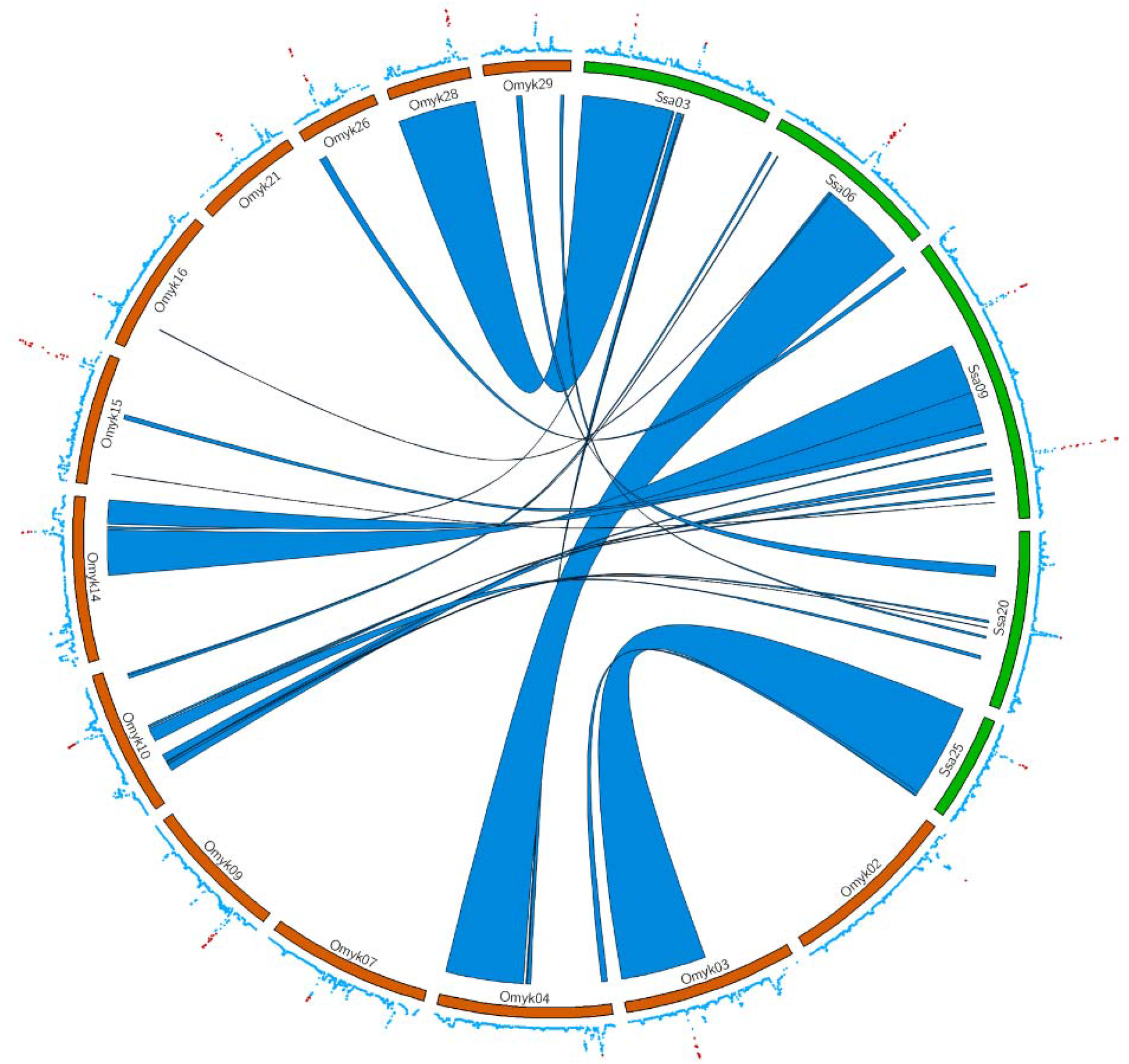
A circos plot of sea lice resistance. The inner ribbons mark syntenic regions between Atlantic salmon (green and labeled *Ssa*) and rainbow trout (orange and labeled *Omyk*) chromosomes. Values from the wssGBLUP analysis are plotted on the outer ring, with significant associations plotted in red (values ≥ 1).

We determined 15 orthogroups were shared between both species (Table S1), which we classified according to gene ontology annotations^55^. One the most interesting groups is orthogroup 12 which contained lysophosphatidic acid receptor 2-like (LPA_2_) of Atlantic salmon and rainbow trout G-protein coupled receptor 12-like and an uncharacterized protein. This orthogroup shares the same GO categories (GO: 0004930, GO: 0007186, GO: 0016021, GO: 0070915, GO: 0007165, GO: 0016020) related to the receptor signaling pathway associated with protein G. The activation of LPA_2_ participates in multiple biological processes, such as cytoskeleton modification via actin fiber formation^56^ and have a role in the activation of related adhesion focal tyrosine kinase (RAFTK)^57^, which in turn participates like a stimulating factor for monocytes and macrophages^58^. In orthogroup 13, we identified *dual specificity protein phosphatase 10-like* (*dust10*) in Atlantic salmon and *dual specificity protein phosphatase 8* (*dust8*) in rainbow trout. These genes have similar annotations in both species, which have the function of inactivating p38^59^ within the MAPK cascade^60^.

## Conclusion

The GWAS performed here for Atlantic salmon and rainbow trout made it possible to compare the genetic basis of sea lice resistance in both species. We found novel information about the resistance of Atlantic salmon and rainbow trout to sea lice, which suggests that there might be a response mediated by leukocytes, and at the same time, the cytoskeleton to promote cell mobility and repair of the wound. The analysis of orthologous proteins provided few characterized proteins, therefore, further investigations of these species are needed to better annotate genes and generate advances in the elucidation of genetics behind resistance to *Caligus rogercresseyi* and other important biologically and economic important traits. Although we did not find common genes explaining resistance between species, we found potential functional genes that can be classified under similar mechanisms. These results suggest that it is possible that similar mechanisms regulate resistance between Atlantic salmon and rainbow trout. Our results provide further knowledge to help establish better control and treatment measures for one of the most important parasitic diseases affecting salmon and trout aquaculture.

## Material and methods

### Rainbow trout

A total of 2,643 rainbow trout were sampled for this study. The fish originated from 105 maternal, 2012 year-class, full-sib families, and belonged to a broodstock population of Aguas Claras S.A company. The fish were separated into three different ponds so that each family was equally represented in each pond. The sea lice infestation was initiated with 105,600 copepodites, on average, an infestation pressure of 40 copepods/fish (produced *in vitro* from ovigerous females). The infestation consisted of depositing the copepodites in each test pond, stopping the flow of water and keeping the pond in darkness for a period of 6 hours. On the sixth day after infestation, parasite counting was performed and caudal fins were sampled for genetic analysis. All the salmon are euthanized and fins were examined for parasite count using a stereoscopic magnifying glass. Wet body weight was recorded for each animal at the end of the challenge.

### Atlantic salmon

A total of 2,628 Atlantic salmon smolts belonging to 118 maternal full-sib families from a 2010 year-class of Salmones Chaicas, X Región, Chile, were challenged with *C. rogercresseyi*. The fish were PIT-tagged (Passive Integrated Transponder), acclimated and distributed into three ponds as described in previous studies^5,8^. Infestation with the parasite was carried out using 13 to 24 copepods per fish, stopping the flow of water for 6 hours after the infestation. The challenge lasted 6 days, then the fish were euthanized and the sea lice were counted on all of the fins. A sample of tail fin was taken for genetic analysis and the wet body weight of each fish was measured.

### Genotyping

Genomic DNA was extracted from the caudal fin of each challenged fish using the DNeasy Blood & Kit tissue kit (Qiagen), following the manufacturer’s instructions. The Atlantic salmon samples were genotyped using an Affymetrix® 50K Axiom® myDesign™ Genotyping Array designed by AquaInnovo and the University of Chile^61^, and the rainbow trout samples were genotyped with a 57K SNP array developed by the USDA^13^.

Quality control of the genotypes was carried out in PLINK^62^. SNPs with a call rate <= 0.95, a major allele frequency (MAF) < 0.05 and those that were not in Hardy-Weinberg equilibrium (p < 1*x*10^-6^) were discarded. Individuals were filtered if they had a call rate <= 0.95. All the SNPs and fish that passed quality control, were used for downstream analysis.

### Genomic Association Analysis

Resistance to *C. rogercresseyi* was defined as follows, according to Ødegård *et al.* (2014)^9^:

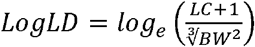

Where LD is the Caligus density defined as the Caligus count (LC) on each fish at the end of the experimental challenge divided by the cube root of the body weight of the fish on the same day (BW) squared, is an approximation of the surface of the skin of each fish. The logarithm of LD was used as it has an approximately normal distribution.

Single step genomic BLUP (ssGBLUP) and wide single step genomic BLUP with two iterations (wssGBLUP)^63^ was used to identify associations between SNPs and resistance to *Caligus rogercresseyi*, using the BLUPF90 family of programs^64^. Both approaches use the combination of a genomic and pedigree matrix. Genotype and pedigree information was used to generate the kinship matrix H^65^ with the following equation:

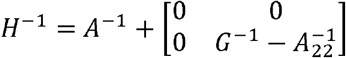

Where *A*^-1^ is the inverse relationship matrix, for all the animals, constructed from the pedigree, 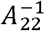 is the inverse of the pedigree matrix produced from genotyped animals, and *G*^-1^ is the reverse matrix of genomic relationship. The SNPs were weighted with equal value and assigned the constant 1 to perform the ssGBLUP method. For the wssGBLUP method, the markers used weights estimated by the previous method. The association analysis for both traits were performed using the following mixed linear model y = **X**b + **Z**a + e, where **y** is the vector of phenotypic values (LogLD); **b** is the fixed effects vector (tank); **a** is the vector for random effects considering the structure of covariance between individuals established by matrix **H**; and **e** is the vector for the random residuals; **X** and **Z** are the incidence matrices for fixed and individual effects respectively.

To identify the regions of the genome associated with the traits analyzed, we identified windows of 20 adjacent SNPs where 1% or more of the phenotypic variance was explained, similar to Neto *et al* (2019)^66^. The cumulative percentage of variance explained for each trait was visualized using a Manhattan plot in R ^67^.

### Genome comparison

The rainbow trout (GCF_002163495.1)^68^ and Atlantic salmon (GCF_000233375.1)^17^ genomes were downloaded from the NCBI and subset for chromosomes associated with sea lice resistance was downloaded using samtools^69^. Synteny between the chromosomes was identified by aligning the sequences using the program Symap v3.4^70^. Circos^71^ was used to plot the relationships between rainbow trout and Atlantic salmon chromosomes and to plot sea lice resistance associations to their respective locations.

### Candidate Genes

The flanking sequences surrounding SNPs associated with sea lice resistance were aligned to the most recent reference genomes of rainbow trout and Atlantic salmon using BLASTn^72^. The sequence was saved in FASTA format. BLASTx was then used to identify coding sequences for proteins in these associated windows. Blast2Go^73^ was used in parallel with the FASTA file to identify proteins and select them by function.

For both species, the reference genome of *Danio rerio* (GenBank Assembly Accession: GCA_000002035.4) was used to annotate proteins that were not characterized in the rainbow trout or Atlantic salmon reference genomes. To identify orthologous proteins/genes between species, the OrthoFinder^74^ program was used with the FASTA sequences obtained with BLASTx.

## Ethics approval and consent to participate

All the experimental challenges were approved by the Comité Institucional de Cuidado y Uso de Animales of the Universidad de Chile (Certificate N 17,041-VET-UCH).

## Consent for publication

Not applicable

## Availability of data and material

Atlantic salmon phenotype and genotype data are available at Figshare (10.6084/m9.figshare.7676147). Rainbow trout phenotypic and genotype data are available in the same repository (https://figshare.com/s/5219597a19f23873fda3).

## Authors’ contributions

PC assessed the analyses and wrote the initial version of the manuscript. AB contributed with results interpretation discussion and writing. KC contributed with genome comparison analysis, writing and discussion. LNB and KC managed samples, performed DNA extraction and performed the quality control of genotypes. JPL contributed with the study design. JMY conceived the study, contributed to results interpretation with a discussion. All authors approved the manuscript.

## Acknowledgments

The current work was funded by Period: 2016-2019, Funding Agency: Research Council United Kingdom and CONICYT, Chile. Project Number: MR/N026144/1 PI: UK: Ross Houston; Chile: José M. Yáñez Co-researcher: John Hickey Institutions: The Roslin Institute - The University of Edinburgh; University of Chile; Aquainnovo

